# Cadmium Disrupts Blood-Testis Barrier (BTB) via An Oxidative Stress-Dependent Autophagy in Prepubertal Rats

**DOI:** 10.1101/2024.06.04.597310

**Authors:** Yonghong Man, Bingxue Du, Jiaolong Huang, Yu Sun, Yunhao Liu, Ling Zhang

## Abstract

Blood testis barrier (BTB) is an important target of cadmium (Cd) toxicology, but the mechanism underlying the Cd-induced impairment of BTB function remains fully elucidated. The mammalian target of rapamycin (mTOR) complex 2 (mTOR2) regulates BTB function via its effects on cellular junctions and the cytoskeleton of Sertoli cells. In this study, we investigated whether mTOR2 was involved in the effects of Cd exposure on BTB integrity. A Cd exposure model *in vivo* was established in prepubertal male rats using a single intraperitoneal injection of Cadmium Chloride (CdCl_2_). A Cd exposure model of Sertoli cell was established using a CdCl_2_-treated TM4 cell line. The Acute cadmium exposure decreased the activity of mTOR2 signaling and adhesin proteins which is linked to the induction of oxidative stress-induced autophagy. In the presence of CdCl_2_, mTOR, the catalytic subunit of the mTOR2 complex, exhibits a reduction in levels of phosphorylation, accompanied by decreased adhesin proteins and Rictor, the key component of the mTOR2 complex. CdCl_2_ treatment also drives a process of oxidative stress-induced autophagy, evidenced by alterations in cellular markers for oxidative stress and autophagy. Pharmaceutical inhibition of oxidative stress and/or autophagy alleviates the alternations in mTOR2 signaling and adhesin proteins upon CdCl_2_ treatment in TM4, a Sertoli cell line. This work is the first to examine the effects of cadmium exposure on rictor/mTOR2 signaling pathways. Our results suggest that Cadmium might exert testicular toxicology via the perturbation in mTOR2 signaling, which can be associated with the cellular stress-related protolysis in Sertoli cells.

## Introduction

Cadmium (Cd), one of the major heavy metals in environmental pollutants, is well-implicated in reproductive disorders [1]. The epidemiological observational studies describe an obscure relationship between Cd exposure and male reproductive function [2]. In contrast, the experimental studies, both *in vivo* and *in vitro*, provided significantevidence for the toxic effects of Cd exposure on reproductive function, including endocrine secretion, spermatogenesis, semen production, and consequently, fertility [3]. The compromised reproductive function is tightly associated with an impairment of seminiferous tubules, which accommodate sustentacular or Sertoli cells, a specific type of Cd-sensitive cell in the testis [4,5]. The versatility of Sertoli cells involves phagocytotic activity, direct connections with differential germ cells, and maintaining a crucial spermatogenesis environment by the formation of a physical barrier, which is known as the blood-testis barrier (BTB) [6,7].

An integral BTB divides the seminiferous epithelium into the basal and the apical (adluminal) compartments with specialized junctions between adjacent Sertoli cells [8,9]. Among these junctions are tight junction and basal ectoplasmic specializations, which form barriers by strands of trans-membrane proteins, such as occludin and E-cadherin [8]. With a terminal end facing the cytoplasm, these trans-membrane proteins establish an association with the actin cytoskeleton via binding proteins (e.g. ZO-1) [10]. A dynamic actin cytoskeleton facilitates the migration of preleptotene spermatocytes from the basal to the adluminal compartment, which occurs during the early stage of spermatogenesis [11]. This transition depends on both junction restructuring and actin cytoskeletal reorganization, a process named BTB dynamics [11]. Decades of biochemical characterizations have defined several relationships between BTB dynamics and regulatory signalings, such as the rapamycin-insensitive companion of mTOR2 [12].

Rictor is a recently identified regulator of BTB function [12]. The stage-specifical expression of Rictor in seminiferous epithelium exhibits a positive correlation with the status of the BTB dynamics [12]. Genetic ablation of Rictor results in the loss of BTB function, highlighted by a perturbance of tight junction, changes in actin organization, and abnormality of protein distribution [13]. Moreover, as a scaffolding protein, Rictor interacts with multiple subunits (e.g. mTOR) to form mTOR complex 2 (mTOR2)[14]. This large complex provides a signaling activity of serine/threonine kinase[14]. And the known substrates of the signaling complex mainly include AGC kinases (Oh W.J., 2011), such as Akt (also known as protein kinase B, PKB)[15]. Thus, the physiological functions of BTB dependent on the homeostasis of both the integrity and dynamics of BTB. Although previous studies have revealed an impairment of BTB function upon cadmium exposure, whether the mTOR2 signaling is involved in this testicular toxicity remains fully elucidated.

Given that Rictor/mTOR2 orchestrates the integrity and dynamics of BTB, we investigate whether cadmium exerts deleterious effects on this signaling complex. Strikingly, we found that Cadmium exposure altered the activities of mTOR2 signaling, both *in vivo and in vitro*, which was accompanied by reduced levels of adhesin proteins (occludin, E-cadherin, ZO-1). Here, we identified oxidative stress-induced autophagy as an important process responsible for these alternations. Mechanistically, CdCl_2_ treatment led to an induction of oxidative stress and enhanced autophagy flux. Pharmaceutical inhibitors decreased the levels of cellular markers for oxidative stress (HO1) and autophagy flux (LC3, p62), respectively. These interventions alleviated the alternations in BTB components after CdCl_2_ treatment. Thus, we showed that cadmium might potentiate a disturbance of BTB functions via Rictor/mTOR2 signaling.

## Materials and methods

### Animals

Sprague-Dawley rats were purchased from the Hubei Experimental Animal Research Center. All animals were first allowed to be accustomed to the new environment for 1 week. They are housed in an animal room that was maintained at 23±1°C and 40±30% relative humidity with air ventilation at a speed of 13 times per hour and a 12-hour light-dark cycle. Rodent Diet standard chow (radiation sterilized) and drinking water (tap water, via automatic water supply system) were provided ad libitum. All animal experiments were performed in compliance with the Guidelines for the regulations approved by the Animal Protection and Experimental Ethics Committee of Wuhan University of Science and Technology (Approval No. 20210305, Wuhan, China).

### Cadmium chloride (CdCl_2_) treatment

CdCl_2_ was purchased from Sigma-Aldrich (St. Louis, MO), and prepared in phosphate-buffered saline (10 mM sodium phosphate and 0.15M NaCl, pH 7.4, at 22°C, PBS) at a concentration of 100mM. The stock solution was then divided into small sterile tubes and stored at -20°C before treatment.

Rats were administrated with CdCl_2_ (0.5, 1, 2mg.kg^-1^, intraperitoneal (i.p.) injection) for 24 hours. The rats of the blank control group were administrated with sterilized phosphate-buffered saline (PBS). One testis of rats was collected for western blot analysis, and the other was processed for histological analysis. In another group of rats, both testes that underwent BTB integrity assay were collected for immunofluorescence analysis as described below.

### BTB integrity assay

BTB integrity *in vivo* was assessed based on its ability to block the entry of small fluorescence probes (e.g., EZ-Link Sulfo-NHS-LC-Biotin) from the basal to the adluminal compartment. In short, rats were anesthetized using Chloral hydrate (350 mg.kg^-1^ bw, i.p.). Thereafter, a small incision (≈1cm) was made on the scrotum to expose the testis. 1mg of Biobin-conjugated EZ-Link Sulfo-NHS-LC (NO.21335, ThermoFisher) dissolved freshly in 100μl PBS (containing 1mM CaCl_2_) was injected under the tunica albuginea into the basal compartment of testis using a micro syringe (Gaoge, Shanghai, China). The other testis was injected with 100μl PBS (containing 1mM CaC_2_) as control. The testes were then collected in Petri dishes with PBS and allowed to rest in an incubator at 37°C for 30min. The testes were thereafter immersed in the Modified Davidson's fixative (mDF) for 6h at room temperature, followed by 4% paraformaldehyde (PFA) in PBS (pH 7.4) (Servicebio Ltd., China), for 18h at 4°C. Paraffin sections (5-μm-thick) of testis were obtained thereafter and processed conventionally. After dewaxing and rehydration, the sections of the testis were incubated with Alexa Fluor@568 streptavidin (2ug/ml) in PBS for 1 hour at room temperature. These stained sections were examined by an FV1000 confocal laser scanning biological microscope (Olympus, JAN).

### Histological analysis

Paraffin sections (5-μm-thick) of testis were conventionally processed, and stained using hematoxylin and eosin (H&E) according to the standard histochemical protocols. The sections were examined with a BX51 Olympus microscope (Olympus Corp., Melville, NY; Center Valley, PA), and photographs were captured with a ProgRes C14 camera (Jenoptik Laser, Jena, Germany).

### Immunofluorescence assay of paraffin sections

For sections immunofluorescence, Paraffin sections (5-μm-thick) testis underwent antigen restoration with citrate antigen retrieval solution (pH6.0). The retrieval sections were blocked in 5% goad serum in PBS (v/v, blocking solution), followed by overnight incubation at 4°C with primary antibodies diluted in the blocking solution. Thereafter, the sections were washed in PBS and incubated with Alexa Fluor– conjugated secondary antibodies (Abcam; green fluorescence, Alexa Fluor 488) at 1:250 dilution with blocking solution for 1h at room temperature. Fluorescence images were captured using FV1000 confocal laser scanning biological microscope (Olympus, JAN).

### Cell culture

The Sertoli cell line TM4 (ATCC number: CRL-1715) was purchased from the Procell Life Science & Technology Co., Ltd. (Wuhan, China), and is authenticated by Short tandem repeat (STR) profiling analysis. Cells were cultured in a growth medium, consisting of a combination of Dulbecco’s modified Eagle’s medium (DMEM) and Ham’s F-12 at a ratio of 1:1, with 17.5mM D-Glucose, 15.1 mM 4-(2-hydroxyethyl)-1-piperazine ethane sulfonic acid (HEPES), 2.5mM glutamine, 0.5 mM sodium pyruvate (Dalian Meilun Biotechnology Co., Dalian, China), supplemented with 5% v/v fetal bovine serum (FBS) (Tianhang Bio, Hangzhou, China) and 1% (v/v) penicillin/streptomycin sulfate cocktail at 37°C in a humidified atmosphere containing 5% CO_2_. When cells reached 70% confluence, they were subjected to incubation under the conditions described below.

### Cell experiment conditions

TM4 cells were seeded in plates at a density of 3×10^4^/cm^2^. The ratio of medium volume to plate surface area is 0.3125ml/cm^2^, and the incubation medium consists of DMEM/F12 (1:1) supplemented with 1% FBS. Then, the cells were allowed to reattach and settle for 12h in an incubator at 37°C. Subsequently, the cells were subjected to incubation for 24h under the following experimental conditions: (a) culture medium alone; (b) CdCl_2_ (5μM); (c) CdCl_2_ (10μM); (d) N-acetylcysteine (NAC, 3mM); (e) NAC (3mM) + CdCl_2_ (10μM); (f) Chloroquine (CQ, 50uM); (g) CQ (50uM) + CdCl_2_ (10μM). Both NAC and CQ were added into the culture medium 1h before the CdCl_2_ treatment.

### Protein isolation

The total protein extraction of the testis or cell culture was prepared using a cell lysis buffer (20mM Tris, 150mM NaCl, 1% Triton X-100, and inhibitor cocktails), supplemented with 1mM Phenylmethanesulfonylfluoride (PMSF). The lysate was then centrifuged at a speed of 12000g at 4°C for 5min, and the supernatant was collected. The protein concentration of the supernatant was determined using a Bicinchoninic Acid (BCA) Protein Assay Kit (Dalian Meilun Biotechnology Co., Dalian, China) by absorption at 562nm. The protein samples prepared were either analyzed immediately or stored at -80°C for future analysis.

### Immunoblotting analysis

The protein samples, mixed with a 6×loading buffer (Beyotime Institute of Biotechnology, Haimen, China), were heated at 95°C for 5min. Equal amounts of protein were separated by SDS-PAGE and transferred to polyvinylidene fluoride membranes (Immobilon-P, Millipore, Bedford, MA). Membranes were blocked with PBS plus 0.1% Tween 20 (PBST) and 5% nonfat dry milk. Membranes were then incubated with primary antibody diluted in 5% milk-PBST at 4°C overnight. Membranes were subsequently washed in PBST and incubated with horseradish peroxidase-conjugated secondary antibodies. Protein detection was performed using enhanced chemiluminescence with the G: BOX Chemi XRQ (Syngene, USA) detection system. Analysis of band intensity was performed using ImageJ software. Antibodies used for immunoblot analysis are listed in Table 1.

**Table 1.** Antibodies used for different experiments in this report.

| **Antibody** | **Host**<br>**species** | **Vendor** | **Catalog**<br>**number** | **Applications**<br>**(s)/dilutions (s)** |
| --- | --- | --- | --- | --- |
| SQSTM1/p62 | Rabbit | CST | 39749 | IB (1:1000), IF (1:100) |
| Atg5 | Rabbit | CST | 12994 | IB (1:1000) |
| Atg7 | Rabbit | CST | 8558 | IB (1:1000) |
| Phospho-Akt<br>(Ser473) | Rabbit | CST | 4060 | IB (1:2000) |
| LC3 | Rabbit | ProteinTech | 14600-1-AP | IB (1:2500) |
| Occludin | Mouse | ProteinTech | 66378-1-Ig | IB (1:10000) |
| ZO-1 | Rabbit | ProteinTech | 21773-1-AP | IB (1:5000), IF (1:200) |
| E-cadherin | Rabbit | ProteinTech | 20874-1-AP | IB (1:10000) |
| Phospho-<br>mTOR<br>(Ser2448) | Rabbit | ProteinTech | 5536 | IB (1:1000) |
| GAPDH | Mouse | ProteinTech | 60004-1-Ig | IB (1:50000) |
| HO-1 | Rabbit | ProteinTech | 10701-1-AP | IB (1:3000) |
| Rictor | Rabbit | ProteinTech | 27248-1-AP | IB (1:4000) |

### Immunofluorescence assay of cell culture

TM4 cells cultured on coverslips were fixed in 4% paraformaldehyde in PBS (w/v) for 10 min and permeabilized in Immunostaining Permeabilization Solution with Triton X-100 (Beyotime Ltd., China) for 5min at room temperature. Cells were then blocked by 5% goad serum in PBS (v/v; blocking solution), followed by overnight incubation at 4°C with primary antibodies diluted in blocking solution: mouse anti-p62 (Proteintech; 1:100), Thereafter, cells were washed with PBS and incubated with Alexa Fluor–conjugated secondary antibodies (Abcam; red fluorescence, Alexa Fluor 555) at 1:250 dilution with blocking solution for 1h at room temperature. Cells were then washed and mounted with Antifade Mounting Medium with 4-,6-diamidino-2-phenylindole (DAPI; Beyotime Ltd. China). Fluorescence images were captured using an Eclipse Ci-L Fluorescence Microscope (Nikon, Japan). For monodansylcadaverine (MDC) staining, cells were only incubated with MDC (Beyotime Ltd., China) diluted in a culture medium according to the manufacturer`s instructions. Fluorescence images were captured using a ZOE Fluorescent Cell Imager (Bio-Rad, USA).

### Statistical Analysis

Experiments were performed in duplicate or triplicate and repeated three times with similar results. Reliable data meet the condition SD/mean<10%. Statistical differences between groups were determined using the independent, two-tailed Student’s t-test. Experimental data are presented as mean±SD (standard deviation). Differences were considered significant when the P value was less than 0.05.

## Results

### Administration of CdCl_2_ changed the construction of seminiferous tubules and barrier function of BTB in rats

Previous studies have shown the ability of cadmium to provoke testicular injury, including histological and biomolecular alterations[16,17]. To determine the dosimetric profile of CdCl_2_ treatment on the testis in prepubertal rats, an *in vivo* model of acute cadmium exposure was established. Using Hematoxylin and Eosin staining, the seminiferous tubules from the group of rats (2mg.kg-1 i.p) exhibited an alternation in the cytoarchitecture of epithelium, highlighted by the heterochromatic and detached germ cells in the adluminal compartment (Figure 1A). However, the basal lamina remained the integrity of the structure. Note that the histologic examination also showed a sign of hyperemia, providing evidence for cadmium-induced vascular injury (Figure 1A, Figure S1). To further examine whether the histological alterations correlated with BTB injury, an in vivo BTB integrity assay was used. The transition of Biobin-conjugated EZ-Link Sulfo-NHS-LC from the basal to the adluminal compartment (Figure 1B) showed a disruption of BTB barrier function in the group of rats (2mg.kg-1 i.p) at 24h after CdCl_2_ treatment. In contrast, this transition of the probe was not observed in other groups of rats (Figure 1B). Cadmium, therefore, exhibited testicular toxicity by altering the construction of seminiferous tubules and BTB barrier function in a dose-dependent manner.

**Figure 1:**
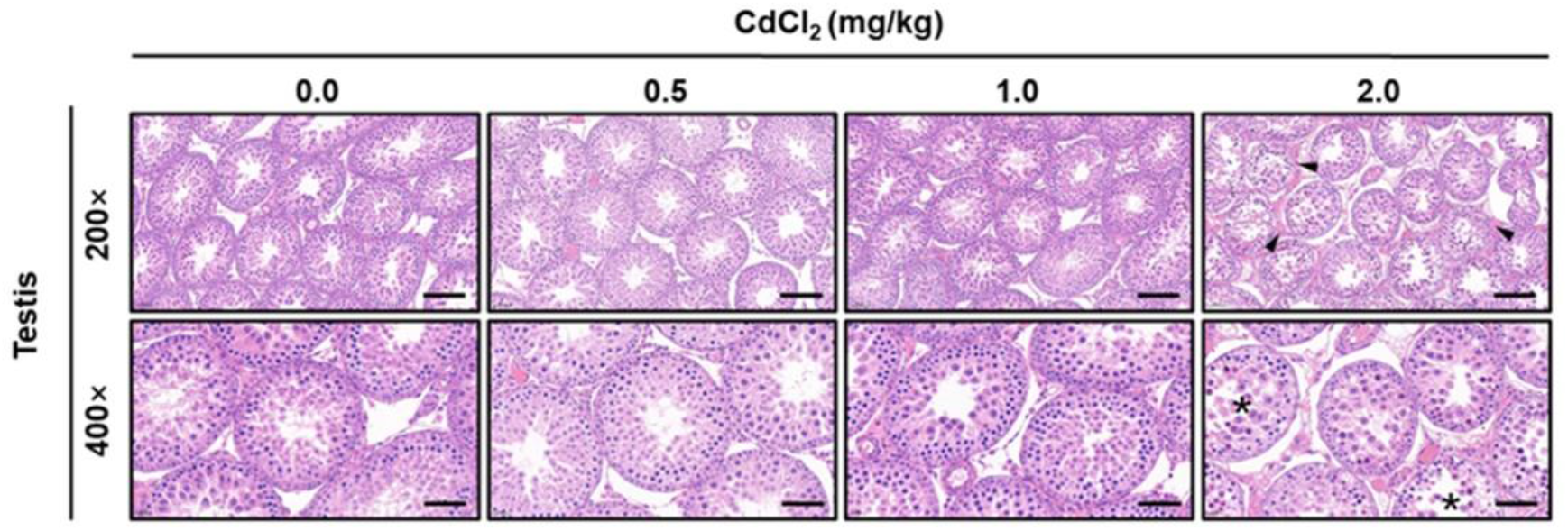
Administration of CdCl_2_ disrupts the BTB integrity in vivo. Prepubertal rats were administrated with either PBS or CdCl_2_ (0.5, 1.0, 2.0mg.kg-1, intraperitoneal (i.p.) injection). Testes were then collected 24h after CdCl_2_ injection for histological microscopy assay.

**Figure 2:**
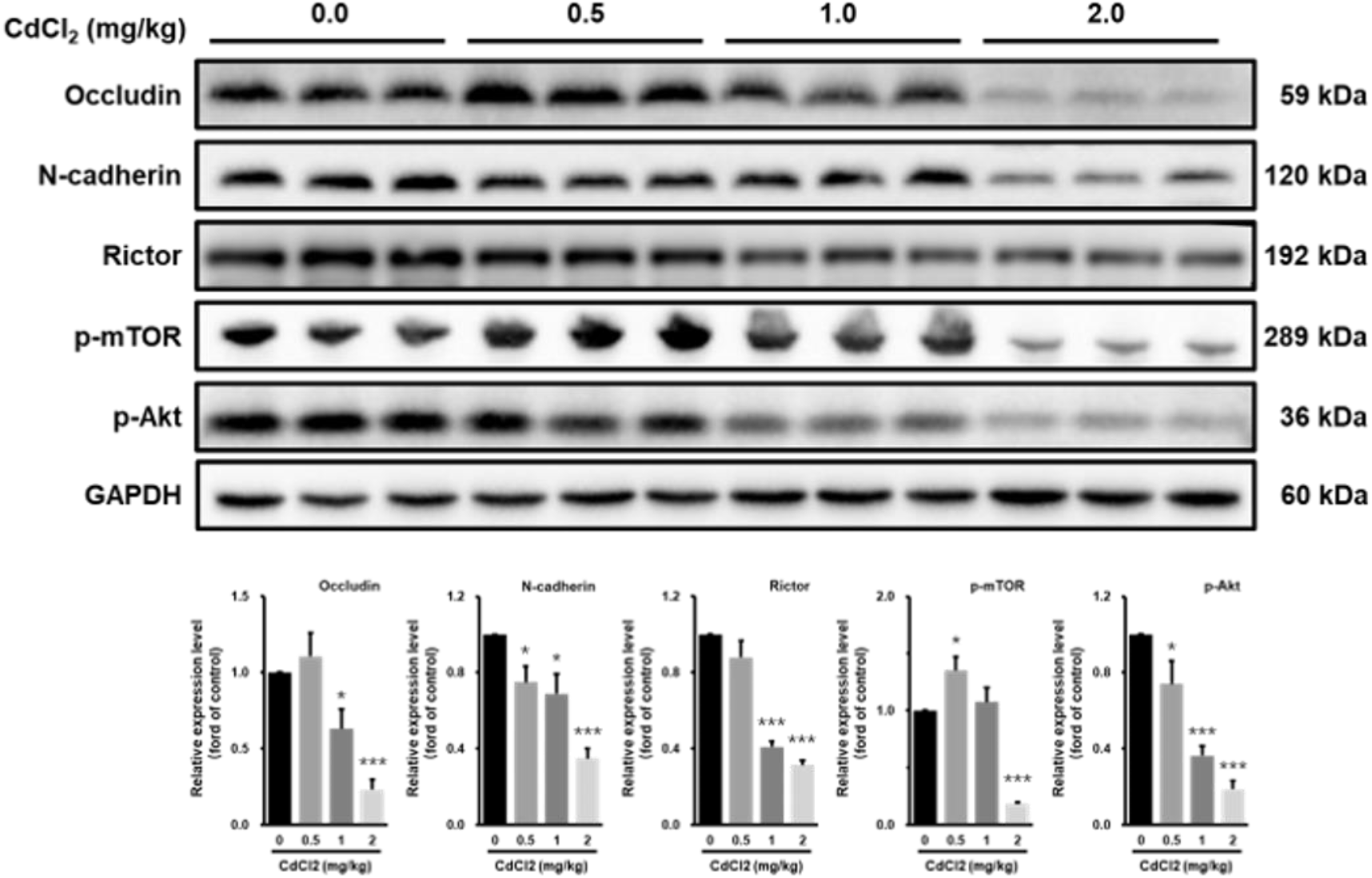
Cadmium exposure changed the adhesin proteins and mTOR2 signaling in vivo.

### Cadmium exposure led to biochemical alternations in BTB components

To determine whether the disruption of the BTB barrier function was related to changes in molecular components involved in BTB, we perform biochemical analysis of the major components of BTB, including both structural proteins (occludin, E-cadherin, and ZO-1) and regulative proteins (Rictor, p-mTOR, and p-Akt). Results showed that acute CdCl_2_ treatment decreased the concentration of occludin and E-cadherin in the testis. The event was more significant in the high-dose group -(CdCl_2_, mg.kg-1, i.p) compared with other groups. The acute CdCl_2_ treatment also led to a decreased level of Rictor, phosphorylated-mTOR (p-mTOR), and phosphorylated-Akt (p-Akt) in a dose-dependent manner. To further validate the destructive effects of cadmium on the tight junction, an immunofluorescence assay was used. The fluorescence staining for ZO-1, another component of tight junction, disappeared from the adjacent Sertoli cells near the basal lamina in CdCl_2_-treated the high-dose group -(CdCl_2_, mg.kg-1, i.p). These data demonstrated that the cadmium-induced BTB disruption was related to an abnormality in the levels of both structural and regulative proteins in the testis.

### The CdCl_2_-induced alternations in BTB components were related to the oxidative stress

Oxidative stress is often implicated in cadmium toxicology, especially in acute cadmium exposure[18]. Our biochemical data further contextualize this stress by an increased MDA level in the testis after CdCl_2_ administration (Figure S2). Previous studies also provided evidence that cadmium induced oxidative stress across diverse cell types[19,20]. And cadmium also leaded to ZO-1 dislocation and downregulation in the blood-brain barrier (BBB)[21]. Based on these data, we hypothesized that oxidative stress might also contribute to cadmium-induced BTB disruption. We use the TM4 Sertoli cell line to model cadmium exposure *in vitro*. The concentration of CdCl_2_ for treatment was determined based on our previous study (Figure S3). Cadmium treatment increased the levels of cellular HO1 (Figure 3A, Figure S3) MDA (Figure S4), and ROS (Figure 8S) in a time- and concentration-dependent manner. In the TM4 cells which are treated with CdCl_2_ for 24h, the cellular levels of adhesin proteins (E-cadherin, Occludin, ZO-1) were reduced significantly (Figure 3, Figure S5). Similar Cadmium-induced, concentration-dependent reduction was also observed in cellular prospered regulatory proteins (p-Akt, p-mTOR, Rictor). These data mirrored the alternations observed in the rat testis upon CdCl_2_ exposure. Note that the transcripts of structural proteins (Occludin, E-cadherin, ZO-1) in CdCl_2_-treated cells were not changed significantly compared with blank groups (Figure S6, Table S1). To determine the contribution of the oxidative stress in these Cadmium-induced molecular events, an anti-oxidant protocol in vitro was used. The addition of NAC (3mM) to the culture medium 1h before CdCl_2_ treatment decreased cellular levels of HO-1 proteins by 72% compared with the positive group (CdCl_2_, 10μM) at 24h after treatment. This protocol also attenuated the reduction in cellular levels of adhesin proteins and prospered regulatory proteins. Cadmium is, therefore, an inducer of the oxidative stress in TM4, and this CdCl_2_-induced stress was related to the abnormalities in adhesin proteins and mTOR2 signaling, both of which are involved in BTB functions.

**Figure 3:**
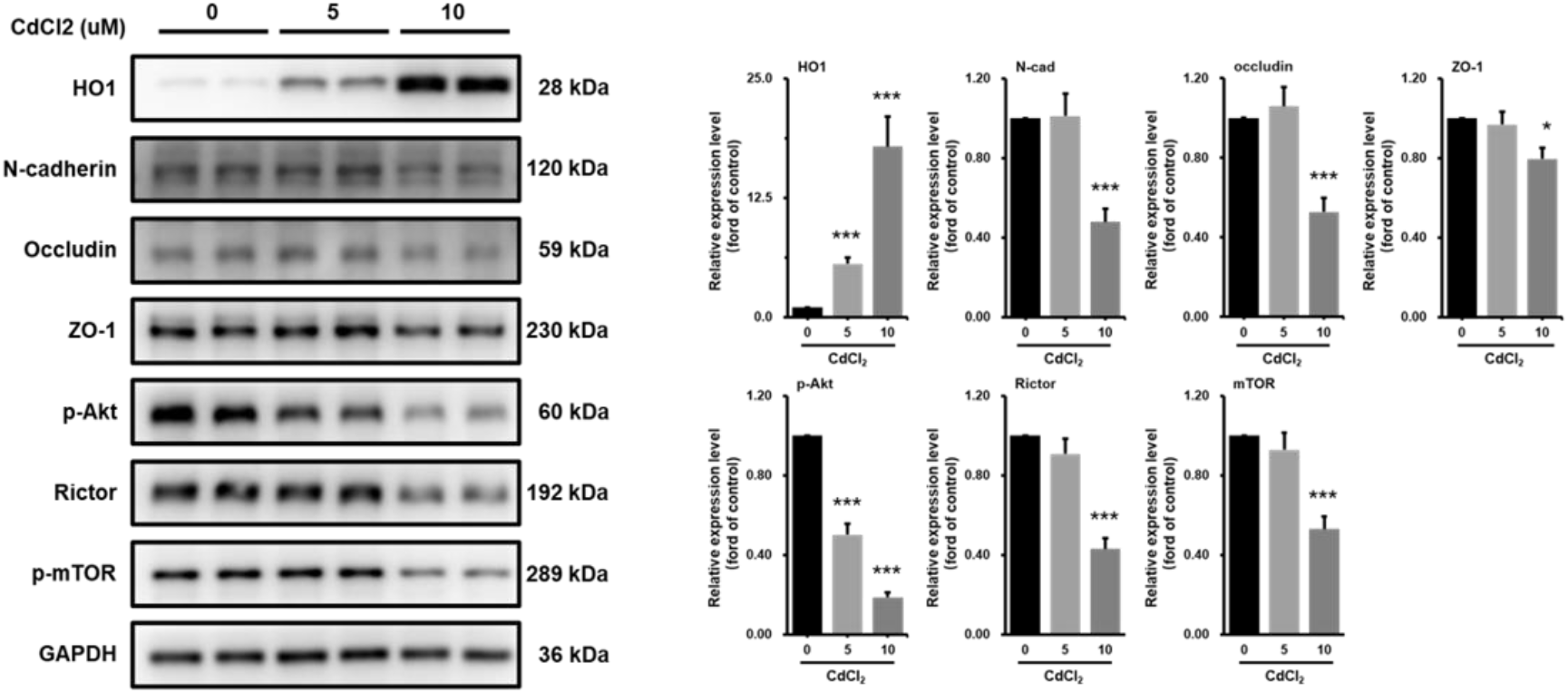
The effects of cadmium on BTB components are related to the oxidative stress.

### The oxidative stress-induced autophagy contributes to the CdCl_2_ toxicity against BTB

The transcriptional and post-translational regulation are two factors that are responsible for protein stability. While the transcripts of adhesin proteins (Occludin, E-cadherin, and ZO-1) are unchanged significantly at 24h after CdCl_2_ (10μM) treatment, we performed biochemical assays of the autophagy. This adaptive response is an essential mechanism for cellular homeostasis. As a major defense against oxidative stress, autophagy removes damaged proteins or organelles via a lysosomal degradation pathway. The administration of CdCl_2_ (0.5, 1, 2mg.kg-1, i.p.) induced a significant increase of LC3B in the testis (Figure S7). CaC_2_ treatment also enhanced the activity of autophagy flux (Figure 4A) and increased the cellular levels of IL3B, p62, Atg5, and Atg7 (Figure 4B, Figure 4C). These proteins are well-established markers for the induction of autophagy flux. The Addition of NAC (3mM) reduced the levels of cellular LC3B and p62 in the presence of CdCl_2_ (Figure 4D), which demonstrate the role of oxidative stress in the CdCl_2_-induced autophagy. Of note, this anti-oxidative protocol did not normalize entirely the cellular p62 level after CdCl_2_ treatment compared with the blank control, which might suggest a different relevance of p62 during cadmium exposure. Nevertheless, the autophagy inhibitor CQ (100uM) alleviated the reduction of adhesion proteins (occludin, E-cadherin) and Rictor compared with the positive control (CdCl_2_, 10μM) (Figure 4E). The suppression of autophagic flux also increased the cellular level of phosphorated mTOR (Figure 4E). These data suggest that oxidative stress-dependent autophagy contributed to the CdCl_2_-induced changes of BTB components in Sertoli cells, including mTOR2 signaling.

**Figure 4:**
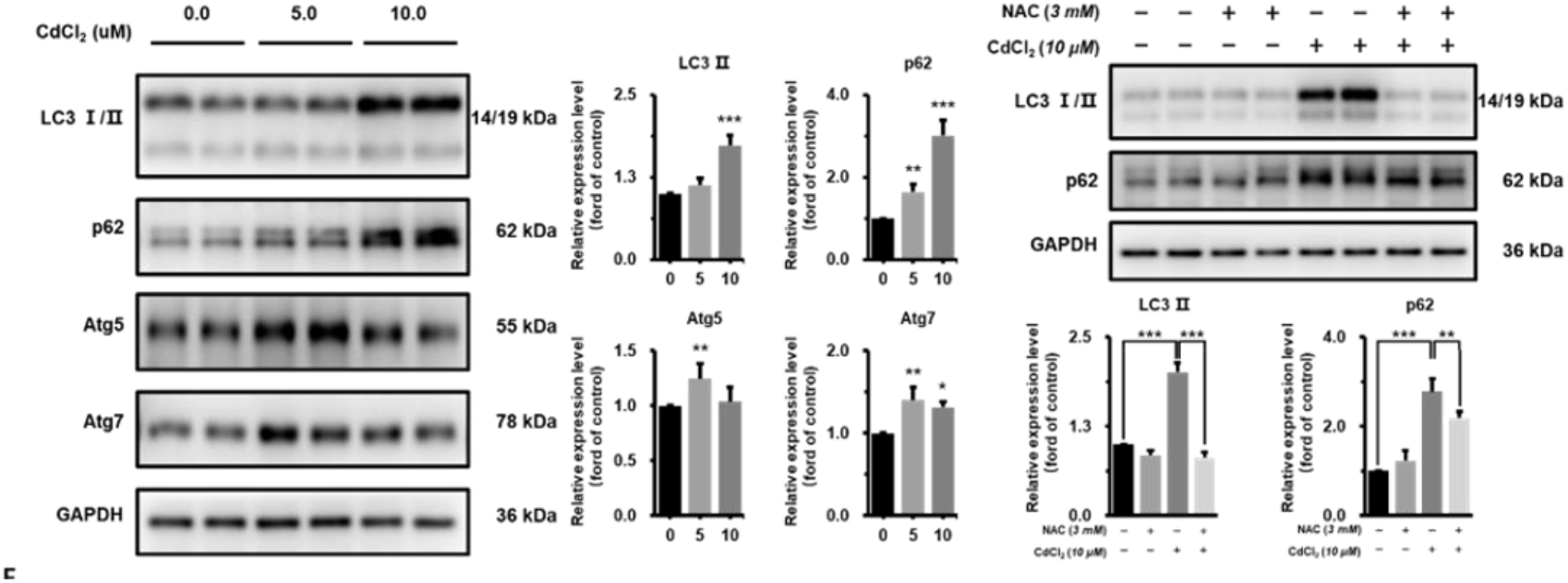
The oxidative stress-induced autophagy was involved in the effects of cadmium on BTB components.

## Discuss

We have identified a role of Oxidative stress-induced autophagy in cadmium toxicology, which might be related to the injury of BTB. Administration of CdCl_2_ resulted in disruption of BTB integrity, evidenced by histologic alternation as well as leakage between basal and adluminal compartments in seminiferous tubules. This Cadmium-induced impairment was associated with abnormalities in both adhesin proteins and regulatory proteins, which are tightly involved in the maintenance of BTB function. Mechanically, oxidative stress and autophagy might be the two critical factors that drive protein degradation upon cadmium exposure in Sertoli cells. These molecular process was linked to alternations in activities of mTOR2 signaling. Therefore, in the presence of Cadmium, the Sertoli cell can exhibit cellular defenses which might also produce interference in the homeostasis of BTB.

BTB is an important target in studies of testicular toxicants. The assessment of BTB integrity provides fundamental evidence for the productive toxicology of a chemical[22–24]. To monitor the BTB integrity, two strategies are employed usually. Both strategies depend on the injection of small fluorescence probes, which can diffuse between basal and adluminal compartments in the presence of BTB injury[12]. The fluorescence probes, including inulin-FITC, and avidin-biotin interaction-based markers, are injected into the jugular vein, or directly into the basal compartment of the testis. To visualize the injected probes, frozen cross-sections of testes were prepared shortly thereafter and examined by fluorescence microscopy. These strategies therefore could complicate the assessment of BTB integrity by excessive operations or the technical limitation of frozen cross sections[12,25]. By contrast, our study allows for the diffusion of fluorescence probes under ex vivo conditions, which cloud simplifies the animal operation protocol. Moreover, the paraffin sections of our study also provided a high-quality visualization of the fluorescence probes, with a more reliable histology structure of the seminiferous epithelium.

Abnormalities in the activity of signaling molecules are involved in BTB damage caused by Cd exposure. Signaling molecules that have been identified to be involved in this injury include Mitogen-activated protein kinase p38 (MAPK p38)[26], Transforming Growth Factor 2 (TGF2)[27], and Transforming Growth Factor 3 (TGF3)[27]. rictor/mTOR2 complex is a newly discovered signaling molecule regulating BTB in recent years[12]. Specific knockdown of rictor in primary Sertoli cell culture disrupts the reconstitution of basal lamina BTB in cell culture *in vitro[12]*. During the process, actin filaments in Sertoli cells undergo rearrangement, accompanied by decreased levels of F-actin and communication junction proteins ( Cx26 and Cx43), as well as redistribution of occludin and ZO-1[12]. These changes suggest that alteration of rictor in Sertoli cells can lead to BTB dysfunction by affecting actin filaments and junction proteins in turn[12]. In the present study, we showed that Cd exposure was able to reduce the level of rictor in rat testis in a dose-dependent manner. The alternation was related to a decrease in the phosphorylation levels both of mTOR and Akt protein. These data suggest that the disruption of the BTB function could be attributed to the dysregulated mTOR2 signaling pathway. This provides new clues to investigate signaling molecular pathways by which Cd affects or damages BTB in mammals.

Previous studies have also shown that oxidative stress-induced impairment of BTB function is associated with changes in junction proteins. Glyphosate (GLY) exposure was able to cause oxidative stress and BTB injury in rat testis, upregulating ROS concentration and downregulating BTB-related proteins in primary Sertoli cells[28]. This injury was shown to be associated with the upregulation of the oxidative stress-related gene NOX1[28]. Exposure to polystyrene microplastics (PS-MPs) induced oxidative stress in mouse testes, accompanied by decreased expression of BTB-related proteins[29]. Methyl parathion (Me-Pa) exposure caused lipid peroxidation and BTB damage in adult male mice[30]. Aflatoxin B1 (AFB1) exposure also caused BTB damage and reduced the expression of BTB-associated junction proteins through oxidative stress in mice[31]. The present study further provides data to support the correlation between oxidative stress and BTB injury. In *in vitro* experiments, CdCl2 treatment induced oxidative stress, evidenced by the increased expression of antioxidant enzymes (HO1). This change was associated with aberrations in junction proteins and regulatory proteins, such as connexins (occludin, E-cadherin, and ZO-1), rictor, p-Akt, and p-mTOR. It is worth noting that these alterations were not normalized entirely by a single pharmacological suppression of oxidative stress. A possible explanation for this event might be that a single NAC was not sufficient for preventing oxidative stress. Indeed, NAC helps to reduce oxidative stress by neutralizing hydrogen peroxide, which is only one of the chemicals contributing to oxidative stress. These data may also implicate the involvement of various stress in BTB injury upon cadmium exposure, such as HRI-responsive mitochondrial stress[32], ER stress[33], and DNA Damage Response[34]. However, these cellular processes share a common activity of inducing autophagy.

Autophagy, a homeostatic cellular process, has undergone an important paradigm shift during the past decades. The emergency of multiple autophagy variants defines a selective removal of cellular components instead of an in-bulk nonselective process. Selective autophagy guarantees the removal of organelles and proteins without provoking other necessary components in the cell[35,36]. The selectivity of autophagy is determined by adaptor protein, and this may suggest a context-dependent activity of autophagy[37]. There is a lack of in-depth studies on the role of autophagy in BTB injury, but relevant studies have demonstrated the correlation between autophagy and adhesion proteins. In porcine mammary epithelial cells, the activation of autophagy coincided with the up-regulation of the expression of tight junction proteins (ZO-1, Claudin-3, and Occludin)[38]; and in a rat model of arterial occlusion/reperfusion (MCAO/R) injury, inhibition of autophagic over-activation was able to promote the expression of tight junction proteins in the blood-brain barrier (occludin, ZO-1, and claudin-5) expression in the blood-brain barrier[39]. In mice, enhanced autophagic activity was able to increase the levels of ZO-1 and Occludin in intestinal epithelial cells, restoring the intestinal mucosal barrier function damaged by sepsis[40]. These studies are controversial regarding the correlation between autophagic activity and adhesion protein expression, which may be related to differences in modeling methods, intervention doses, and observation times. In the *in vitro* experiments, only treatment with a high concentration of CdCl_2_ (10 μM) was able to up-regulate autophagic activity, accompanied by a decrease in the content of connexins (occludin and E-cadherin). By contrast, inhibition of autophagic activity using an autophagy inhibitor (CQ) alleviated reduced occludin and E-cadherin upon Cd exposure. This suggests that autophagy is involved in the degradation of junction proteins; however, inhibition of autophagic activity did not completely restore the level of occludin and E-cadherin to normal levels. This data might suggest that autophagy is not the sole reason for the degradation of junction proteins.

As the key component of the mTOR2 complex, the level of Rictor exhibits a cyclical change in the Sertoli cell during spermatogenesis[12]. The mechanism underlying the diminishment of Rictor around stage VIII in the seminiferous epithelium remains unclear. Moreover, the genetic ablation of Rictor in mice suppresses the levels of actin-binding occludin in the testis[13]. Similarly, our study also observed similar alterations in adhesin proteins upon the reduction of Rictor. Post-translational regulation of protein stability allows for an especially acute alteration in levels, independent of time-consuming transcription[41]. Our data provide a shred of signs for the post-translational regulation of Rictor, evidenced by the different levels of cellular Rictor upon autophagy inhibition. The ubiquitin-proteasome system is another post-translational process responsible for the regulation of protein levels[42]. Note that autophagy inhibition also fails to normalize the levels of Rictor, and this might suggest an involvement of alternative regulations. The precise nature of the interactions between Rictor, autophagy, and ubiquitin-proteasome system warrants further investigation in the future Collectively, our work provides some new insights into the testicular toxicology of cadmium. Acute cadmium exposure exerts deleterious effects on BTB function, which involves the alternations in the activities of the Rictor/mTOR2 signaling pathway. Our data also emphasize the important role of oxidative stress-induced autophagy in this testicular toxicity. The ability of oxidative stress-induced autophagy flux to exert a post-translational regulation on BTB components might suggest a potential therapeutic target in reproductive disorders.

## ACKNOWLEDGEMENT

This work was supported by Hubei Province Higher Education Institutions Young and Middle-aged Innovation Team Project, Environmental Endocrine Disruptors and Population Health [No. T2020003]; Hubei Province Public Health Leading Talents Selection and Cultivation Programme, No. 73, EHTC [2021]; Wuhan Preventive Medicine Special Major Project, Study on the Impact of Environmental Endocrine Disruptor Exposure on Male Reproductive Health and its Mechanism in Wuhan [WY22M01].

